# Genetic Associations with Mathematics Tracking and Persistence in Secondary School

**DOI:** 10.1101/598532

**Authors:** K. Paige Harden, Benjamin W. Domingue, Daniel W. Belsky, Jason D. Boardman, Robert Crosnoe, Margherita Malanchini, Michel Nivard, Elliot M. Tucker-Drob, Kathleen Mullan Harris

## Abstract

Maximizing the flow of students through the science, technology, engineering, and math (STEM) pipeline is important to promoting human capital development and reducing economic inequality^1^. A critical juncture in the STEM pipeline is the highly-cumulative sequence of secondary school math courses^2–5^. Students from disadvantaged schools are less likely to complete advanced math courses, but debate continues about why^6,7^. Here, we address this question using student *polygenic scores*, which are DNA-based indicators of propensity to succeed in education^8^. We integrated genetic and official school transcript data from over 3,000 European-ancestry students from U.S. high schools. We used polygenic scores as a molecular tracer to understand how the flow of students through the high school math pipeline differs in socioeconomically advantaged versus disadvantaged schools. Students with higher education polygenic scores were tracked to more advanced math already at the beginning of high school and persisted in math for more years. Molecular tracer analyses revealed that the dynamics of the math pipeline differed by school advantage. Compared to disadvantaged schools, advantaged schools tracked more students with high polygenic scores into advanced math classes at the start of high school, and they buffered students with low polygenic scores from dropping out of math. Across all schools, even students with exceptional polygenic scores (top 2%) were unlikely to take the most advanced math classes, suggesting substantial room for improvement in the development of potential STEM talent. These results link new molecular genetic discoveries to a common target of educational-policy reforms.

---

Math matters for economic success. American students who take math courses beyond Algebra 2 are more likely to enroll in college, complete a STEM degree ^2–4^, and have better labor market outcomes^5,9,10^. Students from low-income families and schools are unlikely to take advanced math courses in secondary school, which impairs their entry to post-secondary STEM education and ultimately to a STEM career^6,7,11,12^. There are, however, continuing debates about whether the underrepresentation of low-income students in STEM is due to the diminished resources available to their schools and families, or is rather due to those students having lower aptitude or interest in math^7,13–16^. Analyses that statistically control for traditional measures of student interest and ability might lead to biased conclusions about the effects of schools, because these traits can themselves be reciprocally influenced by previous educational experiences ^7^.

Our project addresses the challenge of understanding how student characteristics intersect with school characteristics by using a measure of students’ likelihood to succeed in education that is derived from their DNA. A previous genome-wide association study (GWAS) of 1.1 million people identified hundreds of genetic variants associated with higher educational attainment^8^. These results can be used to calculate an *education polygenic score* (education-PGS)^18–20^, which is a genome-wide composite index of genetic variants associated with completing more years of school. The education-PGS predicts whether or not an individual completes college about as well as his/her family income does^8^. Moreover, unlike traditional measures of student aptitude, individual differences in genetic sequence are fixed at conception and cannot be changed by educational experiences.

Polygenic scores can therefore be used as a molecular tracer to measure flows of students through the STEM pipeline and assess how these flows differ across schools. Just as a radiologist might administer a radioactive tracer to track the flow of blood within the body, researchers can use genetics as a molecular tracer to get a clearer image of how students progress through the twists and turns of a cumulative educational system. Here, we follow the curricular histories of students who vary in their genetic propensity to educational success and who attend secondary schools with varying levels of socioeconomic advantage. This approach offers a novel way of diagnosing the extent to which students who have high genetic propensities for success in education leak out of the STEM pipeline by failing to advance in their mathematics training.

In mapping the flow of students through the secondary school math curriculum, we focus on two dimensions of high school mathematics coursetaking—*tracking* and *persistence*. In some countries (*e.g*., Germany), students are tracked into different types of secondary schools at a discrete number of branch points. The U.S., in contrast, does not have a formal tracking system. Instead, students are offered curricular options that are differentiated by content and difficulty (*e.g*., Pre-Algebra vs. Algebra I vs. Algebra II). Students are informally *tracked* toward final math credentials via their course placement in the first year of secondary school (or earlier)^21^. As subsequent coursetaking hinges on successful completion of pre-requisites and mastery of cumulative content, students’ curricular decisions become strongly path-dependent^21–23^. Students additionally vary after the first year in whether they *persist* in their track throughout secondary school, move to a less-advanced track, or discontinue mathematics training entirely.

First, we sought to validate the education-PGS as a molecular tracer by testing whether it predicted being tracked into a more advanced math class at the beginning of high school and whether it predicted persisting in math for longer. Next, we used the education-PGS to examine differences between schools in the flow of students through the STEM pipeline. Specifically, we focused on the difference between schools that served mainly students from well-educated families versus schools that served mainly students from families with less formal education. Some researchers have proposed that differences between schools in students’ academic achievement are an artifact of school differences in the concentration of students with high genetic propensities toward education^16^. However, students matched on education-PGS differ substantially in their rates of college graduation, depending on where they attended secondary school^24^. Moreover, school characteristics may interact with student genetics by constraining or expanding the range of opportunities for progression through an advanced curriculum. Two students with the same education-PGS might, therefore, differ substantially in their progress through the STEM pipeline, depending on their school characteristics.

## Results

Analyses used genetic and official school transcript data on *N* = 3,635 unrelated adolescents from the National Longitudinal Study of Adolescent to Adult Health (Add Health, see **Methods; Supplementary Figure S1**), who were enrolled in a U.S. high school in 1994-1995. We restricted analyses to European-ancestry participants to prevent inadvertently conflating genetic variation with racial or ethnic background. Previous analyses of national population patterns have revealed a fairly standardized sequence of math coursework, ranging from more basic courses like Pre-Algebra to more advanced courses like Calculus^25,26^. We used this sequence to categorize each participant’s math coursework across four years, based on information obtained from schools, including course catalogs, school information forms, and interviews with school administrators(**Supplementary Table S2**).

At the beginning of secondary school (9^th^ grade, age ~14 years), most students were enrolled in Algebra 1 (51%), but some students were tracked to less advanced (Pre-Algebra or below, 29%) or more advanced (Geometry or above, 20%) courses. A student’s final level of mathematics training was strongly dependent on 9^th^-grade course enrollment: 44% of those enrolled in Geometry or higher in 9^th^-grade ultimately completed Calculus, compared to only 4.2% of those enrolled in Algebra 1 and 1% enrolled in Pre-Algebra or lower level math class. Students with higher polygenic scores were more likely to be tracked into more advanced math courses in 9^th^ grade (Figure 1A, *b* = 0.583, *SE* = .034, *p* = 3.41 × 10^−64^, **Supplementary Table S3**).

**Figure 1.**
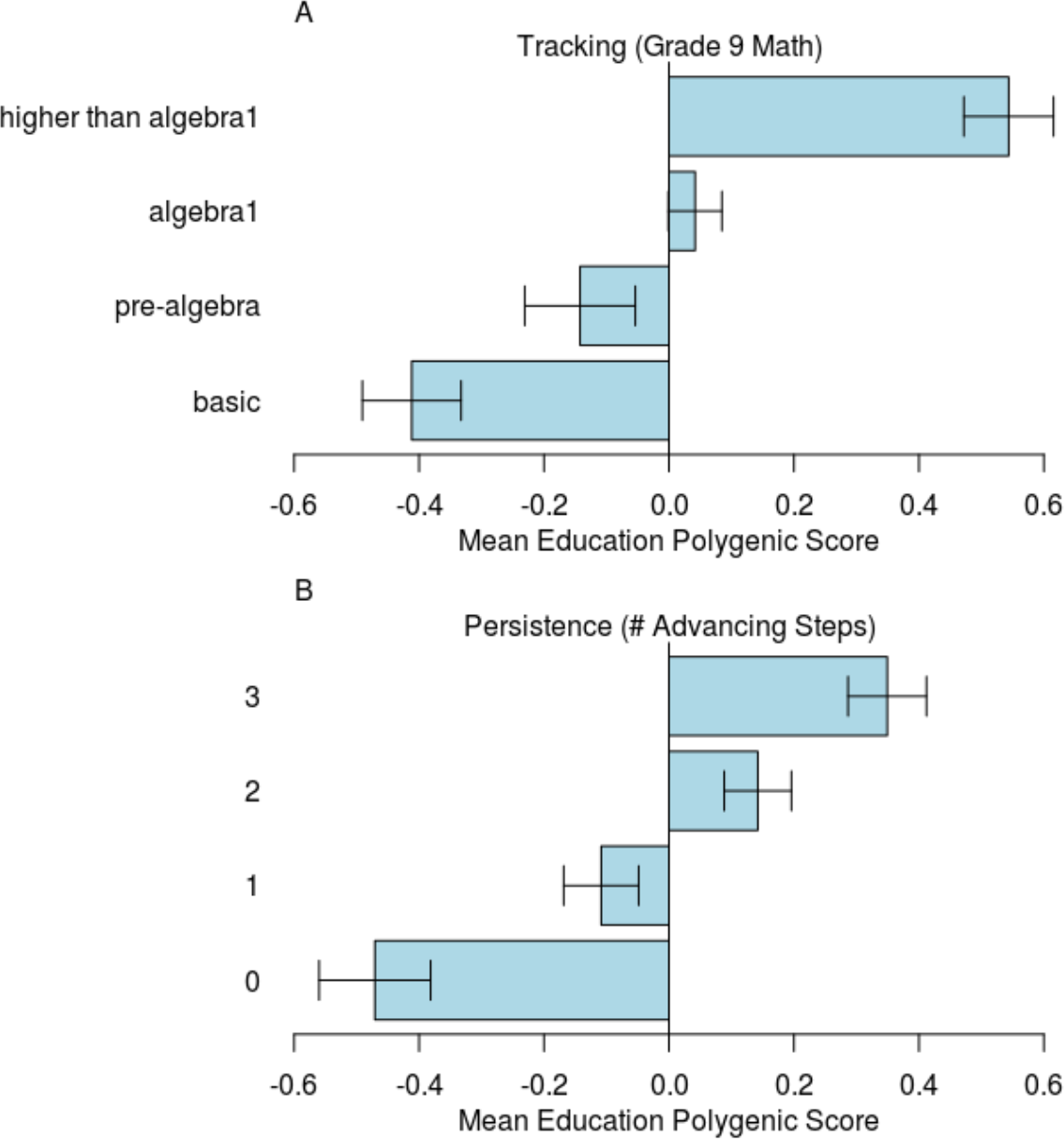
Students with higher education-associated polygenic scores are tracked to more advanced math and persist for longer in math. Error bars represent 95% confidence intervals around the mean.

Add Health participants with a higher education-PGS more often grew up in high-SES families and attended high-SES schools, as compared to participants with lower polygenic scores^24,27^. These gene-environment correlations raise the possibility that genetic associations with mathematics tracking could be due to clustering of students with higher polygenic scores in environmental contexts that better support math achievement. To address this possibility, we repeated our analysis of tracking in the 9^th^ grade using measures of school-SES and family-SES as covariates (**Methods**, **Supplementary Table S3**). As expected, students from higher-SES families were tracked to more advanced math courses at the beginning of secondary school (*b* = 0.419, *SE* = 0.033, *p* < 2 × 10^−16^), as were students in higher-SES schools (*b* = 0.704, *SE* = 0.257, *p* = .0061). However, including family-SES and school-SES as covariates attenuated the association between the education-PGS and mathematics tracking in the 9^th^-grade only by about 20% (attenuated from *b* = 0.583, *SE* = .034, to *b* = 0.461, *SE* = .036, *p* < 2 × 10^−16^, **Supplementary Table S3**). Note that the association with genetics was roughly comparable in magnitude to the association with family-SES.

As a stronger test of whether the genetic association with mathematics tracking was due to clustering of students with high education-PGS into certain schools, we repeated our analysis of 9^th^-grade tracking yet again, this time using school-fixed-effects regression to compare students to their schoolmates^28^ (**Supplementary Table S3**). Comparing only students who were in Algebra 1 or below, students with higher education-PGS were less likely, compared to their schoolmates, to be placed in a remedial track (Pre-Algebra or lower) than in Algebra 1 (*b* = 0.387, *SE* = .049, *p* = 3.65 × 10^−15^). Similarly, comparing only students who were in Algebra 1 or above, students with higher education-PGS were more likely, compared to their schoolmates, to be placed in an advanced track (Geometry or higher) rather than in Algebra 1 (*b* = 0.587, *SE* = .059, *p* = 3.74 × 10^−23^).

What happens to students after the 9^th^-grade? Participants in this sample attended high school in the mid-1990s, when the average high school graduation requirement in U.S. states was 2.4 years of math coursework^29^. Rates of math drop-out accelerated in later years of secondary school (9^th^-grade: 2.6%, 10^th^-grade: 5.2%, 11^th^-grade: 17.6%, 12^th^-grade: 44.7%; **Supplementary Table S2**). Once students dropped out of math, they tended to remain out of math coursework; only 8% of students enrolled in a math class after a year of no math. We summarized persistence across the four years of transcript follow-up as number of advancing steps in the math coursework sequence, ranging from zero to three. For example, a student who completed Algebra 1, Geometry, and Algebra 2, but who did not take a math course in the 12^th^-grade, took two advancing steps.

Students with higher education-PGS took more advancing steps (Figure 1B; *b* = 0.139, *SE* = .013, *p* < 2 × 10^−16^; **Supplementary Table S4**). We then repeated this analysis using a number of additional covariates. As we observed for tracking, students from higher-SES families and who attended higher-SES schools were more likely to persist in math coursetaking across secondary school (family SES *b* = 0.120, *SE* = .014, *p* < 2 × 10^−16^; school SES *b* = 0.234, *SE* = .010, *p* = .018). But the education-PGS association with persistence was only modestly attenuated after accounting for family- and school-SES covariates (*b* = 0.096, *SE* = .014, *p* = 3.1 × 10^−12^). The next model also included 9^th^-grade course placement as a covariate. Students who were tracked to Pre-Algebra or lower in the 9^th^-grade persisted less in math than those in Algebra 1 (*b* = −0.221, *SE* = .034, *p* = 8.2 × 10^−11^). In contrast, students in more advanced math tracks in 9^th^-grade (Geometry or higher) did not differ from those placed in Algebra 1 (*b* = −0.059, *SE* = .035, *p* = 0.087). Controlling for tracking in the 9^th^-grade, the education-PGS again remained associated with persistence (*b* = 0.087, *SE* = .014, *p* = 7.3 × 10^−10^).

As with tracking, we repeated this analysis yet again using school fixed-effects to compare students to others in their school. Consistent with previous analyses, participants with higher education-PGSs took more advancing steps in their mathematics coursetaking than their schoolmates (*b* = 0.117, *SE* = .014, *p* = 1.69 × 10^−17^; **Supplementary Table S4**).

We next analyzed persistence on a year-by-year basis. As shown in Figure 2, most students were enrolled in math in 9^th^-grade, and their mean education-PGS was the sample mean (*i.e*., zero). Few of these students dropped out of math in 10^th^-grade, but these early drop-outs had a low average education-PGS (less than 0.3 *SD* below the mean). The pace of attrition increased in subsequent years (note growth in size of the red dots), and students who continued to take any math class were an increasingly positively-selected group.

**Figure 2.**
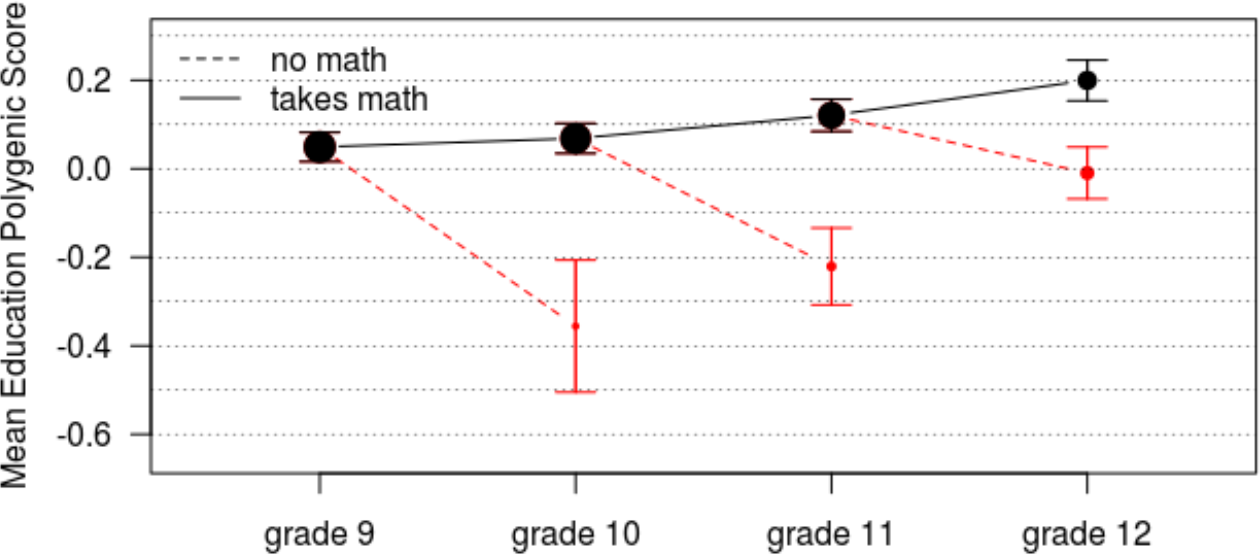
Genetic associations with persistence in math recur year-after-year. Error bars represent 95% confidence intervals. Size of the dots represents number of students enrolled or not enrolled in math in each year.

We considered whether the education-PGS provided any novel information above and beyond what could be observed from students’ performance in math class. It did. This set of analyses focused on students who were enrolled in any math class in the 9^th^-, 10^th^-, and 11^th^-grades, and tested enrollment in any math class in the subsequent year. End-of-year grade point averages (GPAs; on a 4-point scale) in math were obtained from the school transcripts. At every year, students from higher-SES families, students attending higher-SES schools, and students who had higher math GPAs were more likely to enroll in math the subsequent year (**Supplementary Table S5**). After controlling for these covariates, a 1-*SD* increase in the education-PGS was still associated with 1.26 times greater odds of taking a math class in 10^th^-grade (95% CI = 1.06 – 1.51), 1.15 times greater odds in 11^th^-grade (95% CI = 1.04 – 1.28), and 1.13 times greater odds in the 12^th^-grade (95% CI = 1.04 – 1.23).

Our analyses reveal genetically-stratified flows of students through the mathematics training pipeline. We visualized these flows using a “river plot” (Figure 3) ^30^. In the river plot, participants’ math courses (rows) are plotted by year of secondary education (columns). Courses are ordered from most-advanced at the top of the graph to least-advanced at the bottom. The widths of the rivers (*i.e*., the edges connecting row-column nodes) indicate the number of students moving from one course to another. The color of the rivers represents the average education-PGS for students following a particular path (higher in blue, lower in orange). Overall, these results support the premise that the education-PGS can be used as a molecular tracer to evaluate how students flow through the STEM pipeline in secondary school.

**Figure 3.**
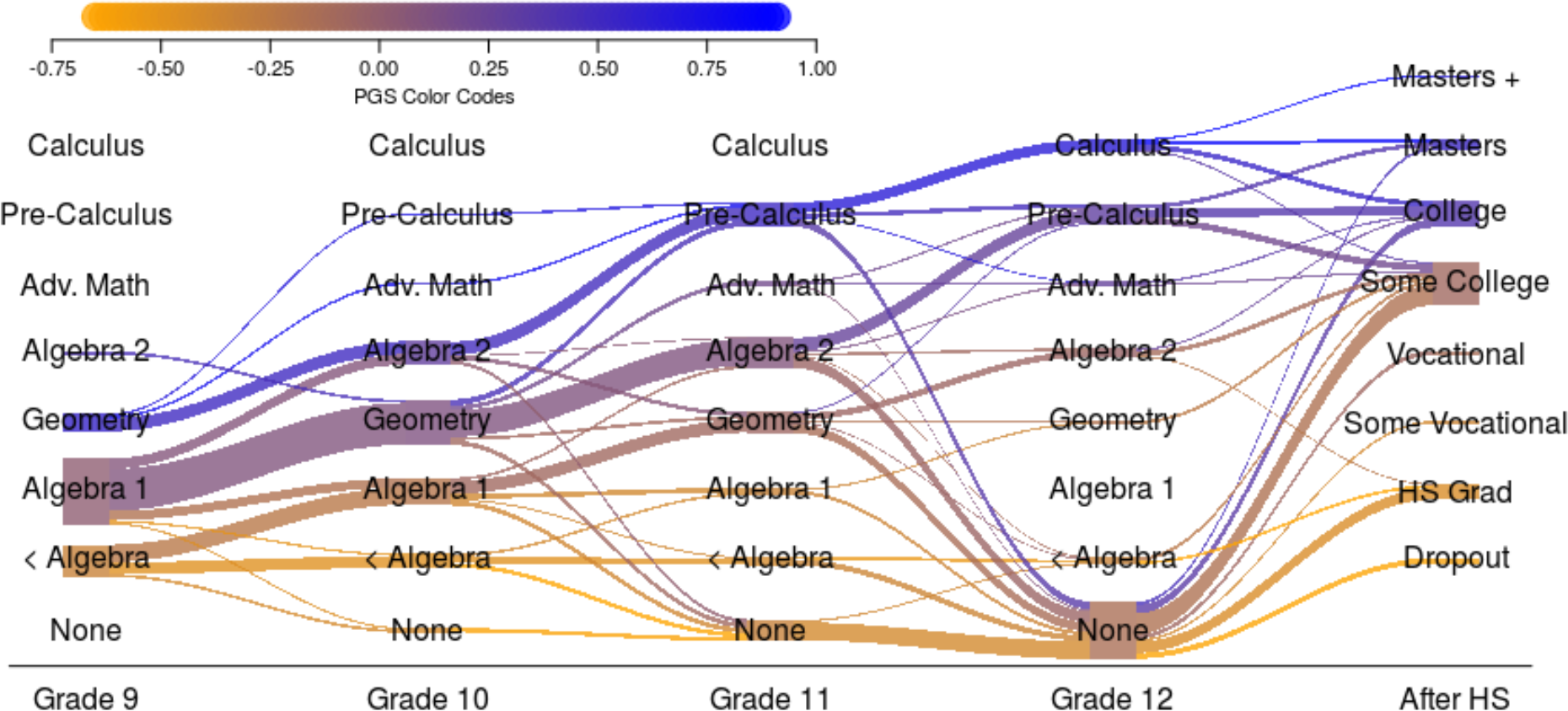
Student DNA can be used to visualize the flow of students through the high school math curriculum. Columns represent year of secondary school; rows represent mathematics course sequence ranging from least to most advanced. Width of the rivers connecting columns proportional to number of students. Shading of rivers represents the average education polygenic score for students in a particular course in a particular year, ranging from low (orange) to high (blue).

We next conducted two analyses of how STEM pipeline dynamics varied by school advantage. First, we tested if the genetic association with tracking differed between high- and low-SES schools using cumulative link models with product terms to capture interactions between school-SES and the education PGS. Students had higher returns to their genetic propensities for educational attainment in higher-status schools (Figure 4A): Higher education-PGS predicted 9^th^-grade tracking more strongly among students in higher-status schools than in lower-status schools (interaction *b* = 0.59, *p* =0.014; **Supplementary Table S3**). A student with an education-PGS of +1 (top 16^th^ percentile) who is in a high-status school has a 33.1% probability of being tracked to Geometry in the 9^th^-grade (note horizontal gray line in Figure 4A). In order to have the same probability of being placed in Geometry, a student in a low-status school would need to have an education-PGS of +2.0 (top 2%). Robustness analyses using non-parametric LOESS and adjacent category logit models suggested similar patterns (see **Supplementary Figure S2**).

**Figure 4.**
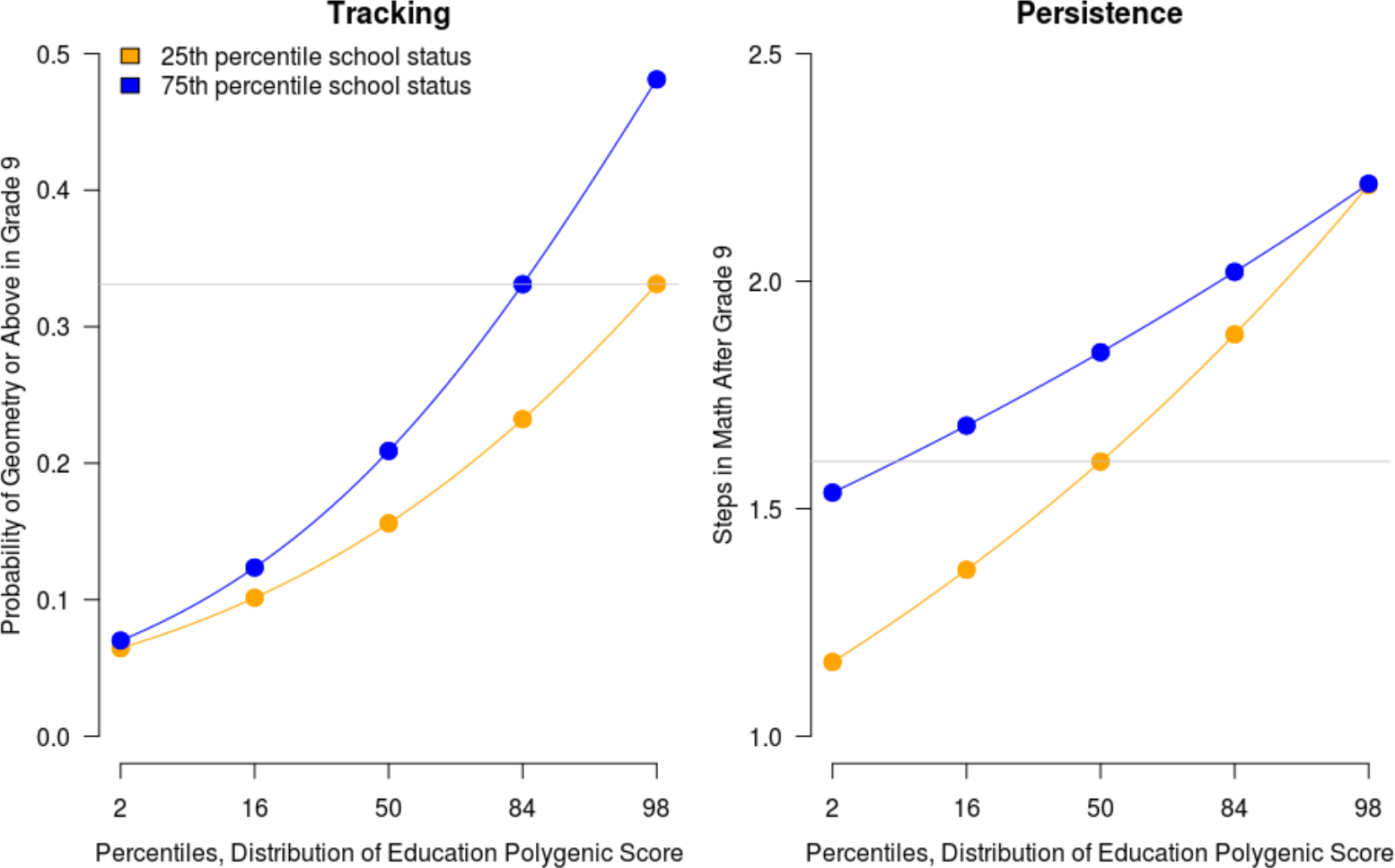
(A) Students with high education-associated polygenic scores are more likely to be tracked into advanced math in advantaged schools than in disadvantaged schools. Fitted probabilities of being tracked to Geometry or higher in the 9^th^ grade, based on cumulative link logit model. School status measured by percent of students whose mother graduated from high school. See Supplementary Table S3 for full model results. (B) **(A) Students with low education-associated polygenic scores persist more in math in advantaged schools than in disadvantaged schools.** Model-implied number of advancing steps from 9^th^- to 12^th^-grade, based on Poisson model. At least 1 year of math beyond the 9^th^ grade was compulsory in most U.S. states. See Supplementary Table S4 for full model results.

Second, we tested the interaction between education-PGS and school-SES in predicting number of advancing steps in math. There was a significant and negative interaction on mathematics persistence, such that low-PGS students were *less* likely to drop out of math in high-status schools than in low-status schools (*b* = −0.304, *SE* = .091, *p* = 8.6 × 10^−04^; **Supplementary Table S4**). The interaction effect was similar when including 9^th^-grade tracking as a covariate (*b* = −0.281, *SE* = .091, *p* = .0036). Figure 4B shows how the number of advancing steps varied as a function of education-PGS in schools at the 0.25 quantile versus 0.75 quantile of school status. High-PGS students persisted about equally in their mathematics training regardless of school status. In contrast, students with an average or low education-PGS were particularly likely to drop out of math in low-SES schools. For example, students attending low-SES schools with an average education-PGS completed 1.6 advancing steps (note gray horizontal line in Figure 4B). However, in high-SES schools, a similarly low level of mathematics persistence is only seen in students at the very low end of the genetic distribution (education-PGS = −1.5, bottom 7%-ile).

Our final analyses focused on school differences in whether or not a student completed Calculus, the most advanced course category in the 9-course sequence. Results from a logistic regression found that school-SES and the education-PGS each predicted taking Calculus, but they did not significantly interact (**Supplementary Table S6)**. Students with an average education-PGS had nearly twice the chances of taking Calculus in a high-SES school (11%) than in a low-SES one (6%). Calculus was rare even among students with exceptional polygenic scores (top 2%, or +2 *SD* above the mean): High-PGS students had a 24% probability of taking Calculus in a low-SES school and a 31% probability in a high-SES school.

## Discussion

In summary, this study used data on student genetics as a molecular tracer to test how the flow of students through the high school math curriculum varied between disadvantaged versus advantaged school contexts. There were three main findings. First, students with higher education polygenic scores tended to enroll in more advanced mathematics tracks in the 9^th^-grade, and they were more likely to persist in these tracks through the end of high school. Second, genetic associations with tracking and persistence were not explained by differences between schools or measured differences in family socioeconomic status. Third, the flows of students through the mathematics pipeline differed between high-SES and low-SES schools.

Specifically, we saw evidence for both cumulative advantage and resource substitution^31,32^. When examining tracking in the 9^th^-grade, the returns to education-linked genetics were higher in high-status schools: Students with high polygenic scores were more likely to be tracked to advanced math classes in higher-status schools. At the same time, students with low polygenic scores were buffered from dropping out of math in high-status schools. The net result of these processes is that students in high-status schools had substantially better math credentials by the end of high school. This difference, we argue, are likely best understood in terms of school differences in resources and practices, rather than in terms of the genetic composition of their student bodies (*cf* ^16^).

It is now well-established that educational attainment is heritable^33^ and can be predicted from an individual’s DNA^34^. What is less well-understood is *how* genetic differences between individuals lead to differences in educational outcomes. In order for genetics research to inform education policy and practice, greater knowledge is needed about the developmental and social processes that connect students’ DNA to their educational outcomes^35^. Indeed, because polygenic scores are largely “black boxes” that aggregate genetic variants with unknown biological functions, some scientists have insisted that genetic discoveries in the area of education have no policy implications whatsoever ^6^.

However, our results directly connect these genetic discoveries to a common target of educational reforms – math coursetaking in secondary school. In the U.S., many states and school districts have increased the number of mathematics courses required for high school graduation^37^, while others have enacted policies designed to push more students into accelerated math tracks^38^, standardize the procedures for deciding how students are tracked^39^, or eliminate tracking altogether^40^. More generally, the U.S. government spends an estimated $3 billion on programs intended to increase STEM degree attainment^41^.

Who benefits the most from these policies and programs? Genetic technology has the potential to provide new insights into this question. For example, one previous study found that a U.K. educational reform, which increased the age at which students could leave school from 15-to 16-years-old, had larger effects on the physical health of people who were at genetic risk for obesity, thus mitigating genetically-associated health disparities^42^. Our results suggest that, in the context of high school math, the answer to “who benefits” might depend on the specific outcome of interest. To our knowledge, there is not enough genetic data available from current cohorts of students to begin to evaluate whether the panoply of local and national policies designed to improve student mathematics education intersects with genetic differences in a similar way, but this is an intriguing direction for future research.

Integrating genetic information in educational research also has the potential to introduce a novel source of data for researchers and policymakers interested in estimating schooling effects on the distribution of student outcomes. Our results suggest that students with the same genetic predisposition can attain very different levels of mathematical training, depending on the school they attend. A persistent methodological problem for using student outcome data to evaluate educational quality is how to separate out the effects of teachers and schools from the effects of out-of-school variables. Family income, in particular, has received considerable attention as a potential confound for measures of school quality: Do students from “good” schools show better performance because their schools provide high-quality instruction or because the schools have low rates of poverty (or because of some dynamic interaction between the two)^43^?

Genetic information about students is as predictive of success in schooling as family income^8^, and genetic differences between students could similarly confound measures of school quality^16,24^. Indeed, one previous study in the U.K. found that value-added measures of teacher quality were correlated with the average education-PGS of students, suggesting that conventional models of educational quality that fail to consider genetic differences between students might lead to biased conclusions,^44^ whereas incorporating data on student genetics might help clarify the impact of schools: In this study, we examined only one school-level characteristic (proportion of students whose mothers graduated high school), so much work remains to be done to identify the characteristics of teachers, schools, and school districts that maximize the outcomes of students *relative* to others who have the same starting point in life with regards to their genetic propensity toward completing education.

Finally, with the caveat that the Add Health data represents an earlier cohort of students, our results suggest that even advantaged school contexts squander an enormous amount of potential human capital. Out of students who both had exceptional polygenic scores (+2 *SD* above the Add Health sample mean or the top 2% of the population distribution) and attended high status high school, about 31% took Calculus by the end of high school, whereas only 24% of students who had the same score and attended low status schools took calculus. This deficit in advanced course-taking among students with exceptional genetic propensities for succeeding in education indicates a tremendous amount of squandered human potential. The pipeline is leaking badly.

We acknowledge several limitations. First, the genetic predictor deployed here captures only a fraction of the genotypic variation relevant for education. Other variants (*e.g*., rare variants^45^) relevant for ultimate educational attainment might not operate through the processes described here. Additionally, this genetic predictor is useful only for understanding individual differences between people of European ancestry, as the validity of education GWAS results has been established only for this segment of the population^8^. The extent to which results will generalize to other populations is uncertain, and none of our results is relevant to understanding disparities between ancestry groups^46^. Until equally well-powered polygenic scores are available for all major ancestry groups represented in U.S. schools, it will not be feasible to use genetic information for policy-relevant decisions.

Second, previous studies have suggested that up to half of the polygenic score association with educational outcomes might operate indirectly, through the parental genotype shaping the quality of the environment provided to children, rather than directly through the biology of the child herself ^47^. Consequently, the association between genotype and math curricular choices might partially operate not through the genetically-influenced characteristics of the student herself, but through the genetically-influenced characteristics of her parents, such as the greater knowledge that college-educated parents have about how to navigate a differentiated curriculum^48^. Disentangling such indirect genetic effects^49^ from genetic effects that operate through the biology of the student will require larger samples of genetic relatives, such as parent-offspring trios^50^. We conducted an initial exploration of this question by comparing siblings raised in the same family (**Supplement**), but the relatively small number of sibling pairs with transcript data available in Add Health limits the definitiveness of our conclusions about the role of indirect genetic effects.

Third, there is limited information on the educational histories of students prior to the 9^th^-grade, but students’ secondary school experiences are, of course, shaped by their previous mathematics skill development and curricular choices. We suspect that the genetic associations with tracking in the 9^th^-grade partially reflects genetic variation in math skills that have been acquired prior to high school; however, the roles of attributes other than math ability, including the constellation of personality and motivational factors referred to as “non-cognitive” skills, are also likely important^51^. We see potential hints of this here, as polygenic scores predict persistence in math *even* after controlling for the student’s math grades in the previous year. Other genetically-influenced traits that are potentially influential for course placement are the student’s attention problems, behavioral and mental health difficulties, academic interests, motivations, and self-concept, and ability to elicit support and investment from adults^52–54^.

As sample sizes for GWAS continue to increase, more and more specific genetic variants associated with complex human phenotypes, like educational attainment, will continue to be identified. There are dangers associated with genetic research being used to justify an overly reductionistic or bio-deterministic account of educational outcomes^55,56^. However, we show here how DNA measures offer new opportunities for educational science. Specifically, we show that genetics can be used to identify leaks in the STEM pipeline and can refine our understanding who is benefitting (and who is not) from advantaged educational contexts. Integration of genetic data into educational research has the potential to accelerate knowledge about which educational contexts maximize success relative to a student’s starting point in life.

## Methods

### Sample

The National Longitudinal Study of Adolescent to Adult Health (Add Health^57^) is a nationally-representative cohort drawn from a probability sample of 80 U.S. high schools and 52 U.S. middle schools (in roughly 90 U.S. communities). Participating schools were representative of all U.S. schools in 1994–95 with respect to region, urban setting, school size, school type, and race or ethnic background.

We constructed our analytic sample as follows (see also **Supplementary Table S1**). At Wave 1, data was collected for *N* = 20,369 respondents. At Wave 3, respondents of the Add Health study, who were then 18-26 years old, were contacted and asked to give signed consent for the release of their official high school transcripts (AHAA^58^). Transcripts were collected regardless of whether the student graduated from high school. At Wave 4, biospecimens were collected for genome-wide genotyping (described in ^27,59^). Of those in the genetic sample, we focused on unrelated respondents of European ancestry, due to the problem of population stratification in diverse samples^46,60^. Transcripts (*N* = 12,032) and genetic samples (*N* = 5,045 of European origin) were collected for partially overlapping subsets of the Wave 1 respondents. Our analytic sample therefore consisted of 3,635 European ancestry respondents with both genotypic and transcript data.

Descriptive statistics are contained in **Supplementary Table S1**. Compared to the full Add Health sample, our analytic sample had higher family SES, higher overall GPAs, and higher rates of postsecondary education. Missing information further reduced sample size in some analyses.

### Measures

#### Polygenic score

Using results from the most recent educational attainment GWAS^8^, we constructed a polygenic score using all SNPs with reported effect sizes that are also in the Add Health genetic dataset and where the effect allele can be reliably matched to the allele reported in the Add Health genetic data. We residualized the PGS on the top 10 principal components of genetic ancestry and then standardized the PGS based on the full set of respondents in the genetic dataset (*N* = 5,045). A similar PGS has been used in previous work^24,27^. Genotyped respondents who were not in the transcript dataset had a mean education-PGS of −0.11, whereas the genotyped respondents with transcript data had a mean education-PGS of 0.04 (**Supplementary Table S1**).

#### Transcript course-taking indicators

Course content information obtained from the schools was used to identify the level of each course on a student’s transcript and to assign it a Classification of Secondary School Courses code. These codes were used to develop an ordinal indicator of the math course sequence, ranging from 0 (no math) to 9 (calculus). These indicators were developed by the AHAA project^58,61^ to be compatible with the 2000 National Assessment of Educational Progress High School Transcript Study^62^ and are based on population patterns of coursetaking as derived from the National Education Longitudinal Study of 1988^63^. The percentages of students enrolled in each level at each year are in **Supplementary Table S2**. For analysis of 9^th^-grade coursetaking, math courses that focus on remedial skills (Basic/Remedial and General) were collapsed, as were math classes beyond Geometry (Algebra 2, Pre-Calculus, Advanced Math, and Calculus).

#### Course grades

Students’ final math course grades at each year were obtained from transcripts and coded on a 0-4 scale (0=*F*, 1=*D*, 2=*C*, 3=*B*, 4=*A*). If a student took the class pass/fail, withdrew, or received an incomplete, then his/her course grade is missing^26^. A cumulative GPA was also computed from these transcript-based grades.

#### Family socioeconomic status

Family-of-origin socioeconomic status (SES) was indexed using the first principal component of parental education, job status, income, and the number of benefits received (loadings were 0.58, 0.43, 0.49, and 0.49 respectively); see ^27^ for additional information on this indicator. The family-SES variable was standardized with respect to the full Wave I sample; the current analytic sample was more advantaged than the full sample (*M* = 0.34, *SD* = 1.16).

#### School socioeconomic status

We used an indicator of school status used previously^24,64^. Add Health administered an in-school survey to all students in participating high schools (*N* = 90,118). This information was used to calculate the percentage of students at each school who report that their mother graduated high school.

#### Analyses

Our statistical models varied as a function of the outcome variable. For non-categorical outcome variables (e.g., number of advancing steps), we fit baseline generalized linear models of the form:

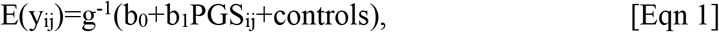

where *i* indexes school, *j* indexes individual, and an appropriate link function *g* is chosen given the distribution of the outcome *y*. For analyses of 9^th^-grade tracking, we fit ordered logistic regressions^65^. For analysis of persistence, we used Poisson regressions. We fit interaction models of the form:

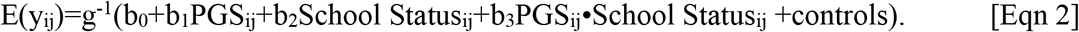

For interaction models, we also included interactions between covariates and the key main effects, so as to guard against spurious findings from specification error^66^. Thus, in models examining interactions between the education-PGS and School Status (as in Figure 4), we also included interactions between PGS and sex, School Status and sex, PGS and birth year, and School Status and birthyear (see **Supplementary Table S3**).

For our ordinal categorical outcomes (*e.g*., tracking in 9^th^-grade), we consider cumulative link models^65^. As used here, cumulative link models assert that:

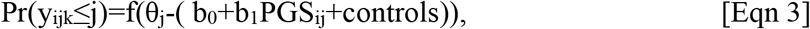

where *k* in [0,1,…*K*] now indexes the category of the outcome *y*. We used a logit link, rendering this model equivalent to the proportional odds model ^67^. One key assumption of this model is that the effect of the predictors does not vary across categories. We therefore also present results from alternative models (*e.g*., adjacent-category logit models) as robustness checks, see **Supplement**.

