## Supplementary Information for "Genetic Associations with Mathematics Tracking and Persistence in Secondary School"

**Supplement**

**A. Descriptives**

Tables S1, S2, and Figure S1 describe sample.

**B. Supporting Tables**

Tables S3-S6 contains estimates on which we report in the main text.

**C. Sensitivity Analyses**

Figure S2 consider non-parametric LOESS and adjacent category logit model (which do not require a homogeneous effect across categories; *c.f.*, Figure 4) as alternatives to the cumulative link models presented in main text.

**D. Results from Family Fixed-Effects Models**

We also conducted exploratory analyses of a sample of *N* = 441 sibling pairs. (One participant per sibling pair was selected to be included in the primary analytic sample). Within-sibling differences in genotype are entirely random, and siblings raised in the same home share many aspects of their home environments. Testing whether siblings who differ in their education-PGS also differ in their subsequent education outcomes is, therefore, a strong test of genetic effects. We used family fixed-effects regression to compare siblings raised in the same nuclear family (*N* = 441 sibling pairs)^1^. As with the school fixed-effects models, we fitted two logistic regression models that focused on deviations from the modal 9^th^-grade track (Algebra 1). The magnitude of the regression coefficients in the family fixed-effects models were similar to the previous analyses (Table S3**)**: When examining students who were in Algebra 1 or lower in the 9^th^-grade, students with higher education-PGS were less likely than their siblings to be placed in a remedial track (Pre-Algebra or lower) than in Algebra 1 (*b* = 0.592, *SE* = .547, *p* = 0.28). Similarly, when examining students who were in Algebra 1 or higher in the 9^th^ grade, students with higher education-PGS were more likely than their siblings to be placed in an advanced track (Geometry or higher) versus Algebra 1 (*b* = 0.473, *SE* = 0.39, *p* = 0.23). However, neither effect was significantly different from zero.

To analyze persistence, we used the same Poisson regression analysis of advancing steps we used in other analyses. In contrast to the results seen for tracking, the magnitude of the regression coefficient for the education-PGS was attenuated relative to the coefficients estimated in previous models (*b* = 0.049, *SE* = 0.078, *p* = 0.53) and not significantly different from zero (Table S4**)**.

Previous analyses of genetic data from trios of parents and offspring have shown that polygenic score associations with educational outcomes might partly reflect indirect genetic effects of the *parental* genotype, which are mediated via environments provided by the parent to all children in the family, *i.e.*, “genetic nurture” ^2,3^. Consistent with this conclusion, the GWAS of educational attainment that was the basis of the education-PGS used in this study found that polygenic effect sizes were attenuated approximately 40% when comparing within-families relative to what was observed when comparing unrelated people ^4^. As the effects of the education-PGS on tracking and persistence were not significantly different from zero, we cannot rule out the possibility that the education-PGS is reflecting the genetic endowment of the students’ parents. This indirect genetic effect could operate through environmental advantages provided by the parents (such as parental knowledge and social capital) rather than a direct effect operating through the students’ own skills and characteristics. The magnitude of the within-family coefficient for tracking, however, was similar to the effect estimated in between-family models, suggesting that these indirect genetic effects might be more relevant for understanding persistence than tracking.

Table S1. Comparison of analytic sample to full Add Health sample.

|  | All (*N*=20369, those with valid school id) | | | Transcript (*N*=12032) | | | Genetic (*N*=5045) | | | Analytic (*N*=3635) | | |
| --- | --- | --- | --- | --- | --- | --- | --- | --- | --- | --- | --- | --- |
|  | *Mean* | *SD* | # NA | *Mean* | *SD* | # NA | *Mean* | *SD* | # NA | *Mean* | *SD* | # NA |
| Cumulative GPA | 2.57 | 0.84 | 8408 | 2.57 | 0.84 | 71 | 2.69 | 0.81 | 1429 | 2.69 | 0.81 | 19 |
| SES | 0.01 | 1.35 | 1392 | 0.11 | 1.29 | 673 | 0.28 | 1.19 | 203 | 0.34 | 1.16 | 125 |
| School Status | 0.48 | 0.15 | 603 | 0.48 | 0.14 | 405 | 0.47 | 0.15 | 193 | 0.47 | 0.14 | 145 |
| Female | 0.5 | 0.5 | 2 | 0.53 | 0.5 | 0 | 0.53 | 0.5 | 0 | 0.54 | 0.5 | 0 |
| Birthyear | 78.83 | 1.73 | 15 | 78.91 | 1.73 | 8 | 78.97 | 1.74 | 1 | 79.02 | 1.75 | 1 |

Table S2. Percentages of students in courses during each year of high school.

|  | Year 1 | Year 2 | Year 3 | Year 4 |
| --- | --- | --- | --- | --- |
| No math | 2.6 | 5.2 | 17.6 | 44.7 |
| Basic/Remedial | 4.3 | 2.6 | 1.5 | 1.1 |
| General | 9.7 | 5.6 | 5.1 | 5.7 |
| Pre-algebra | 12.4 | 3.4 | 1.5 | 0.7 |
| Algebra 1 | 51.3 | 21.4 | 6.6 | 2.3 |
| Geometry | 14.6 | 36.5 | 17.6 | 5.2 |
| Algebra 2 | 4.2 | 19.1 | 24.7 | 8.3 |
| Adv Math | 0.3 | 2.7 | 4.8 | 4.8 |
| Pre-calculus | 0.4 | 3.1 | 19.2 | 17.2 |
| Calculus | 0.1 | 0.2 | 1.2 | 10.0 |

Table S3. Tracking regression coefficients

|  | baseline | baseline plus ses | baseline plus ses.both | baseline.interaction |
| --- | --- | --- | --- | --- |
| basic\|pre-algebra | -1.829 | -1.785 | -1.464 | -1.102 |
| basic\|pre-algebra.se | 0.058 | 0.059 | 0.131 | 0.160 |
| basic\|pre-algebra.pv | 1.16E-217 | 1.82E-201 | 4.92E-29 | 5.26E-12 |
| pre-algebra\|algebra1 | -1.067 | -0.987 | -0.665 | -0.330 |
| pre-algebra\|algebra1.se | 0.051 | 0.052 | 0.128 | 0.158 |
| pre-algebra\|algebra1.pv | 3.49E-98 | 2.71E-81 | 2.01E-07 | 3.62E-02 |
| algebra1\|higher than algebra1 | 1.457 | 1.665 | 1.994 | 2.246 |
| algebra1\|higher than algebra1.se | 0.054 | 0.057 | 0.133 | 0.163 |
| algebra1\|higher than algebra1.pv | 3.67E-159 | 2.11E-185 | 1.34E-50 | 5.88E-43 |
| ea3 | 0.583 | 0.469 | 0.461 | 0.283 |
| ea3.se | 0.034 | 0.035 | 0.036 | 0.124 |
| ea3.pv | 3.41E-64 | 5.22E-40 | 2.37E-38 | 2.21E-02 |
| sexmale | -0.261 | -0.275 | -0.275 | -0.660 |
| sexmale.se | 0.066 | 0.067 | 0.067 | 0.234 |
| sexmale.pv | 7.86E-05 | 3.58E-05 | 3.54E-05 | 4.88E-03 |
| birthyear | 0.088 | 0.088 | 0.088 | 0.278 |
| birthyear.se | 0.019 | 0.019 | 0.019 | 0.067 |
| birthyear.pv | 4.13E-06 | 5.40E-06 | 5.54E-06 | 3.44E-05 |
| sespc.all |  | 0.452 | 0.419 |  |
| sespc.all.se |  | 0.031 | 0.033 |  |
| sespc.all.pv |  | 3.57E-49 | 3.47E-37 |  |
| school.hsgrad |  |  | 0.704 | 1.570 |
| school.hsgrad.se |  |  | 0.257 | 0.320 |
| school.hsgrad.pv |  |  | 6.13E-03 | 9.41E-07 |
| ea3:school.hsgrad |  |  |  | 0.592 |
| ea3:school.hsgrad.se |  |  |  | 0.241 |
| ea3:school.hsgrad.pv |  |  |  | 1.41E-02 |
| ea3:sexmale |  |  |  | -0.049 |
| ea3:sexmale.se |  |  |  | 0.068 |
| ea3:sexmale.pv |  |  |  | 4.71E-01 |
| ea3:birthyear |  |  |  | -0.006 |
| ea3:birthyear.se |  |  |  | 0.020 |
| ea3:birthyear.pv |  |  |  | 7.43E-01 |
| school.hsgrad:sexmale |  |  |  | 0.833 |
| school.hsgrad:sexmale.se |  |  |  | 0.475 |
| school.hsgrad:sexmale.pv |  |  |  | 7.94E-02 |
| school.hsgrad:birthyear |  |  |  | -0.408 |
| school.hsgrad:birthyear.se |  |  |  | 0.138 |
| school.hsgrad:birthyear.pv |  |  |  | 3.13E-03 |
| (Intercept) |  |  |  |  |
| (Intercept).se |  |  |  |  |
| (Intercept).pv |  |  |  |  |
| N | 3367 | 3367 | 3367 | 3367 |
|  | alg1.baseline | alg1.ses | alg1.school | alg1.fam |
| basic\|pre-algebra |  |  |  |  |
| basic\|pre-algebra.se |  |  |  |  |
| basic\|pre-algebra.pv |  |  |  |  |
| pre-algebra\|algebra1 |  |  |  |  |
| pre-algebra\|algebra1.se |  |  |  |  |
| pre-algebra\|algebra1.pv |  |  |  |  |
| algebra1\|higher than algebra1 |  |  |  |  |
| algebra1\|higher than algebra1.se |  |  |  |  |
| algebra1\|higher than algebra1.pv |  |  |  |  |
| ea3 | 0.388 | 0.299 | 0.387 | 0.592 |
| ea3.se | 0.043 | 0.045 | 0.049 | 0.547 |
| ea3.pv | 3.77E-19 | 2.54E-11 | 3.64E-15 | 2.80E-01 |
| sexmale | -0.191 | -0.208 | -0.245 | -2.276 |
| sexmale.se | 0.082 | 0.084 | 0.091 | 0.716 |
| sexmale.pv | 1.98E-02 | 1.30E-02 | 7.18E-03 | 1.48E-03 |
| birthyear | 0.072 | 0.075 | 0.174 | 0.298 |
| birthyear.se | 0.024 | 0.024 | 0.036 | 0.197 |
| birthyear.pv | 2.41E-03 | 1.97E-03 | 1.67E-06 | 1.30E-01 |
| sespc.all |  | 0.380 |  |  |
| sespc.all.se |  | 0.038 |  |  |
| sespc.all.pv |  | 7.48E-24 |  |  |
| school.hsgrad |  |  |  |  |
| school.hsgrad.se |  |  |  |  |
| school.hsgrad.pv |  |  |  |  |
| ea3:school.hsgrad |  |  |  |  |
| ea3:school.hsgrad.se |  |  |  |  |
| ea3:school.hsgrad.pv |  |  |  |  |
| ea3:sexmale |  |  |  |  |
| ea3:sexmale.se |  |  |  |  |
| ea3:sexmale.pv |  |  |  |  |
| ea3:birthyear |  |  |  |  |
| ea3:birthyear.se |  |  |  |  |
| ea3:birthyear.pv |  |  |  |  |
| school.hsgrad:sexmale |  |  |  |  |
| school.hsgrad:sexmale.se |  |  |  |  |
| school.hsgrad:sexmale.pv |  |  |  |  |
| school.hsgrad:birthyear |  |  |  |  |
| school.hsgrad:birthyear.se |  |  |  |  |
| school.hsgrad:birthyear.pv |  |  |  |  |
| (Intercept) | 0.728 | 0.680 | 17.615 | 19.688 |
| (Intercept).se | 0.058 | 0.059 | 783.682 | 11951.104 |
| (Intercept).pv | 1.37E-36 | 6.65E-31 | 9.82E-01 | 9.99E-01 |
| N | 2719 | 2719 | 2719 | 332 |
|  | >alg1.baseline | >alg1.ses | >alg1.school | >alg1.fam |
| basic\|pre-algebra |  |  |  |  |
| basic\|pre-algebra.se |  |  |  |  |
| basic\|pre-algebra.pv |  |  |  |  |
| pre-algebra\|algebra1 |  |  |  |  |
| pre-algebra\|algebra1.se |  |  |  |  |
| pre-algebra\|algebra1.pv |  |  |  |  |
| algebra1\|higher than algebra1 |  |  |  |  |
| algebra1\|higher than algebra1.se |  |  |  |  |
| algebra1\|higher than algebra1.pv |  |  |  |  |
| ea3 | 0.534 | 0.451 | 0.587 | 0.473 |
| ea3.se | 0.051 | 0.052 | 0.059 | 0.390 |
| ea3.pv | 9.12E-26 | 7.22E-18 | 3.74E-23 | 2.26E-01 |
| sexmale | -0.212 | -0.220 | -0.199 | 0.063 |
| sexmale.se | 0.096 | 0.097 | 0.106 | 0.732 |
| sexmale.pv | 2.69E-02 | 2.36E-02 | 6.12E-02 | 9.31E-01 |
| birthyear | 0.079 | 0.076 | 0.106 | -0.223 |
| birthyear.se | 0.028 | 0.028 | 0.042 | 0.243 |
| birthyear.pv | 4.71E-03 | 7.59E-03 | 1.16E-02 | 3.58E-01 |
| sespc.all |  | 0.379 |  |  |
| sespc.all.se |  | 0.051 |  |  |
| sespc.all.pv |  | 1.54E-13 |  |  |
| school.hsgrad |  |  |  |  |
| school.hsgrad.se |  |  |  |  |
| school.hsgrad.pv |  |  |  |  |
| ea3:school.hsgrad |  |  |  |  |
| ea3:school.hsgrad.se |  |  |  |  |
| ea3:school.hsgrad.pv |  |  |  |  |
| ea3:sexmale |  |  |  |  |
| ea3:sexmale.se |  |  |  |  |
| ea3:sexmale.pv |  |  |  |  |
| ea3:birthyear |  |  |  |  |
| ea3:birthyear.se |  |  |  |  |
| ea3:birthyear.pv |  |  |  |  |
| school.hsgrad:sexmale |  |  |  |  |
| school.hsgrad:sexmale.se |  |  |  |  |
| school.hsgrad:sexmale.pv |  |  |  |  |
| school.hsgrad:birthyear |  |  |  |  |
| school.hsgrad:birthyear.se |  |  |  |  |
| school.hsgrad:birthyear.pv |  |  |  |  |
| (Intercept) | -1.081 | -1.292 | -17.642 | -20.765 |
| (Intercept).se | 0.066 | 0.075 | 780.142 | 12496.506 |
| (Intercept).pv | 9.21E-61 | 3.74E-67 | 9.82E-01 | 9.99E-01 |
| N | 2400 | 2400 | 2400 | 334 |

Table S4. Persistence regression coefficients

|  | baseline | baseline plus ses.both | baseline plus ses.both+math9 | baseline.interaction | baseline.interaction.math9control | school | fam |
| --- | --- | --- | --- | --- | --- | --- | --- |
| (Intercept) | 0.541 | 0.383 | 0.466 | 0.255 | 0.313 | 0.533 | 0.928 |
| (Intercept).se | 0.018 | 0.050 | 0.051 | 0.063 | 0.076 | 0.159 | 0.459 |
| (Intercept).pv | 1.10E-197 | 1.17E-14 | 7.80E-20 | 5.08E-05 | 4.35E-05 | 7.90E-04 | 4.31E-02 |
| ea3 | 0.139 | 0.096 | 0.087 | 0.268 | 0.232 | 0.117 | 0.049 |
| ea3.se | 0.013 | 0.014 | 0.014 | 0.048 | 0.052 | 0.014 | 0.078 |
| ea3.pv | 6.87E-26 | 3.07E-12 | 7.25E-10 | 2.58E-08 | 8.80E-06 | 1.69E-17 | 5.31E-01 |
| sexmale | -0.058 | -0.058 | -0.050 | -0.126 | -0.101 | -0.060 | -0.003 |
| sexmale.se | 0.027 | 0.027 | 0.027 | 0.094 | 0.094 | 0.027 | 0.113 |
| sexmale.pv | 3.06E-02 | 3.11E-02 | 6.25E-02 | 1.81E-01 | 2.82E-01 | 2.68E-02 | 9.79E-01 |
| birthyear | -0.008 | -0.009 | -0.012 | 0.005 | -0.009 | 0.002 | -0.044 |
| birthyear.se | 0.008 | 0.008 | 0.008 | 0.027 | 0.027 | 0.011 | 0.033 |
| birthyear.pv | 2.69E-01 | 2.17E-01 | 1.13E-01 | 8.57E-01 | 7.38E-01 | 8.78E-01 | 1.88E-01 |
| sespc.all |  | 0.120 | 0.106 |  |  |  |  |
| sespc.all.se |  | 0.014 | 0.014 |  |  |  |  |
| sespc.all.pv |  | 3.15E-18 | 4.17E-14 |  |  |  |  |
| school.hsgrad |  | 0.234 | 0.210 | 0.613 | 0.633 |  |  |
| school.hsgrad.se |  | 0.099 | 0.099 | 0.125 | 0.151 |  |  |
| school.hsgrad.pv |  | 1.83E-02 | 3.46E-02 | 8.70E-07 | 2.68E-05 |  |  |
| math.start-1 |  |  | -0.221 |  | -0.112 |  |  |
| math.start-1.se |  |  | 0.034 |  | 0.125 |  |  |
| math.start-1.pv |  |  | 8.24E-11 |  | 3.70E-01 |  |  |
| math.start1 |  |  | -0.059 |  | 0.068 |  |  |
| math.start1.se |  |  | 0.035 |  | 0.115 |  |  |
| math.start1.pv |  |  | 8.72E-02 |  | 5.53E-01 |  |  |
| ea3:school.hsgrad |  |  |  | -0.304 | -0.282 |  |  |
| ea3:school.hsgrad.se |  |  |  | 0.091 | 0.097 |  |  |
| ea3:school.hsgrad.pv |  |  |  | 8.60E-04 | 3.58E-03 |  |  |
| ea3:sexmale |  |  |  | 0.008 | 0.010 |  |  |
| ea3:sexmale.se |  |  |  | 0.027 | 0.027 |  |  |
| ea3:sexmale.pv |  |  |  | 7.57E-01 | 7.19E-01 |  |  |
| ea3:birthyear |  |  |  | -0.001 | 0.000 |  |  |
| ea3:birthyear.se |  |  |  | 0.008 | 0.008 |  |  |
| ea3:birthyear.pv |  |  |  | 9.40E-01 | 9.77E-01 |  |  |
| school.hsgrad:sexmale |  |  |  | 0.135 | 0.106 |  |  |
| school.hsgrad:sexmale.se |  |  |  | 0.185 | 0.185 |  |  |
| school.hsgrad:sexmale.pv |  |  |  | 4.63E-01 | 5.67E-01 |  |  |
| school.hsgrad:birthyear |  |  |  | -0.027 | -0.007 |  |  |
| school.hsgrad:birthyear.se |  |  |  | 0.053 | 0.053 |  |  |
| school.hsgrad:birthyear.pv |  |  |  | 0.607 | 0.902 |  |  |
| ea3:math.start-1 |  |  |  |  | 2.62E-02 |  |  |
| ea3:math.start-1.se |  |  |  |  | 0.034 |  |  |
| ea3:math.start-1.pv |  |  |  |  | 0.444 |  |  |
| ea3:math.start1 |  |  |  |  | -3.12E-03 |  |  |
| ea3:math.start1.se |  |  |  |  | 0.036 |  |  |
| ea3:math.start1.pv |  |  |  |  | 0.931 |  |  |
| school.hsgrad:math.start-1 |  |  |  |  | -3.07E-01 |  |  |
| school.hsgrad:math.start-1.se |  |  |  |  | 0.262 |  |  |
| school.hsgrad:math.start-1.pv |  |  |  |  | 0.241 |  |  |
| school.hsgrad:math.start1 |  |  |  |  | -1.71E-01 |  |  |
| school.hsgrad:math.start1.se |  |  |  |  | 0.214 |  |  |
| school.hsgrad:math.start1.pv |  |  |  |  | 0.422 |  |  |
| N | 3367 | 3367 | 3367 | 3367 | 3367 | 3367 | 441 |

Table S5. Year-by-year persistence models

|  | Grade 9 to Grade 10 | | Grade 10 to Grade 11 | | Grade 11 to Grade 12 | |
| --- | --- | --- | --- | --- | --- | --- |
| (Intercept) | 1.930 | 1.219 | 0.988 | 0.296 | 0.054 | -0.507 |
| (Intercept).se | 0.309 | 0.352 | 0.181 | 0.206 | 0.142 | 0.164 |
| (Intercept).pv | 4.12E-10 | 5.38E-04 | 4.39E-08 | 1.51E-01 | 7.03E-01 | 1.99E-03 |
| ea3 | 0.288 | 0.234 | 0.220 | 0.143 | 0.167 | 0.123 |
| ea3.se | 0.085 | 0.090 | 0.049 | 0.051 | 0.040 | 0.041 |
| ea3.pv | 6.81E-04 | 9.20E-03 | 8.28E-06 | 5.17E-03 | 2.69E-05 | 2.58E-03 |
| sespc.all | 0.199 | 0.104 | 0.233 | 0.182 | 0.154 | 0.109 |
| sespc.all.se | 0.068 | 0.074 | 0.043 | 0.044 | 0.038 | 0.039 |
| sespc.all.pv | 3.57E-03 | 1.58E-01 | 4.65E-08 | 3.85E-05 | 5.22E-05 | 5.18E-03 |
| school.hsgrad | 2.931 | 2.717 | 1.650 | 1.654 | 0.652 | 0.720 |
| school.hsgrad.se | 0.682 | 0.699 | 0.379 | 0.387 | 0.286 | 0.290 |
| school.hsgrad.pv | 1.71E-05 | 1.02E-04 | 1.36E-05 | 1.96E-05 | 2.24E-02 | 1.32E-02 |
| sexmale | -0.302 | -0.183 | -0.236 | -0.117 | 0.161 | 0.245 |
| sexmale.se | 0.160 | 0.168 | 0.094 | 0.096 | 0.076 | 0.078 |
| sexmale.pv | 5.97E-02 | 2.76E-01 | 1.18E-02 | 2.25E-01 | 3.49E-02 | 1.77E-03 |
| birthyear | -0.050 | -0.062 | 0.021 | 0.017 | -0.023 | -0.033 |
| birthyear.se | 0.046 | 0.048 | 0.027 | 0.027 | 0.022 | 0.022 |
| birthyear.pv | 2.76E-01 | 1.94E-01 | 4.25E-01 | 5.27E-01 | 2.77E-01 | 1.35E-01 |
| gpa |  | 0.370 |  | 0.302 |  | 0.231 |
| gpa.se |  | 0.076 |  | 0.043 |  | 0.034 |
| gpa.pv |  | 1.16E-06 |  | 2.12E-12 |  | 1.70E-11 |
| N | 3676 | 3652 | 3573 | 3539 | 3087 | 3050 |

Table S6. Highest math credential models.

|  | Highest math is at least Algebra 2 | Highest math is calculus |
| --- | --- | --- |
| (Intercept) | -0.041 | -3.534 |
| (Intercept).se | 0.191 | 0.304 |
| (Intercept).pv | 8.31E-01 | 3.04E-31 |
| ea3 | 0.466 | 0.897 |
| ea3.se | 0.155 | 0.226 |
| ea3.pv | 2.70E-03 | 7.17E-05 |
| school.hsgrad | 1.867 | 2.406 |
| school.hsgrad.se | 0.398 | 0.567 |
| school.hsgrad.pv | 2.65E-06 | 2.17E-05 |
| sexmale | -0.350 | -0.662 |
| sexmale.se | 0.274 | 0.422 |
| sexmale.pv | 2.02E-01 | 1.17E-01 |
| birthyear | 0.232 | 0.367 |
| birthyear.se | 0.078 | 0.120 |
| birthyear.pv | 3.15E-03 | 2.24E-03 |
| ea3:school.hsgrad | 0.313 | -0.385 |
| ea3:school.hsgrad.se | 0.317 | 0.399 |
| ea3:school.hsgrad.pv | 3.24E-01 | 3.35E-01 |
| ea3:sexmale | -0.029 | 0.015 |
| ea3:sexmale.se | 0.084 | 0.124 |
| ea3:sexmale.pv | 7.30E-01 | 9.02E-01 |
| ea3:birthyear | 0.031 | -0.014 |
| ea3:birthyear.se | 0.024 | 0.035 |
| ea3:birthyear.pv | 1.96E-01 | 6.99E-01 |
| school.hsgrad:sexmale | -0.010 | 1.148 |
| school.hsgrad:sexmale.se | 0.571 | 0.769 |
| school.hsgrad:sexmale.pv | 9.85E-01 | 1.35E-01 |
| school.hsgrad:birthyear | -0.454 | -0.550 |
| school.hsgrad:birthyear.se | 0.166 | 0.222 |
| school.hsgrad:birthyear.pv | 6.15E-03 | 1.34E-02 |
| N | 3367 | 3367 |

### Figure S1. Construction of analytic sample.

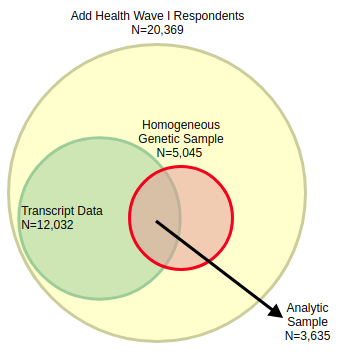

Figure S2. Robustness analyses for math course placement in first year of postsecondary school. Each column considers the probability of placement in a math course relative to a course (or set of courses) immediately below as a function of EA PGS. The top row shows LOESS estimated probabilities while the bottom shows probabilities from a logit model (conditional on being female and of mean birthyear) for students in schools in the top and bottom half of the school SES distribution. Orange and blue represent relatively high and low sttus schools (respectively). The bottom panels represent fitted probabilities based on logistic regression models similar to the cumulative link models discussed in the main text.

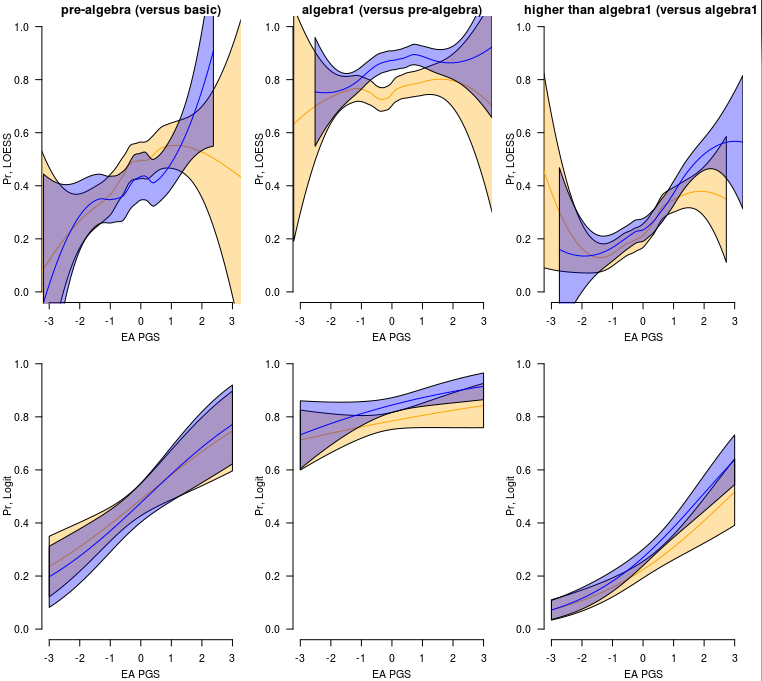
